# DNA methylation in *Verticillium dahliae* requires only one of three putative DNA methyltransferases, yet is dispensable for growth, development and virulence

**DOI:** 10.1101/2020.08.26.268789

**Authors:** H. Martin Kramer, David E. Cook, Grardy C.M. van den Berg, Michael F. Seidl, Bart P.H.J. Thomma

**Affiliations:** Laboratory of Phytopathology, Wageningen University and Research, Droevendaalsesteeg 1, 6708 PB Wageningen, the Netherlands; Department of Plant Pathology, Kansas State University, 1712 Claflin Road, Manhattan, Kansas 66506, USA; Theoretical Biology & Bioinformatics, Department of Biology, Utrecht University, Utrecht, The Netherlands; University of Cologne, Institute for Plant Sciences, Cluster of Excellence on Plant Sciences (CEPLAS), 50674 Cologne, Germany

## Abstract

DNA methylation is an important epigenetic control mechanism that in many fungi is restricted to genomic regions containing transposons. Two DNA methyltransferases, Dim2 and Dnmt5, are known to perform methylation at cytosines in fungi. While most ascomycete fungi encode both Dim2 and Dnmt5, only few functional studies have been performed in species containing both. In this study, we report functional analysis of both *Dim2* and *Dnmt5* in the plant pathogenic fungus *Verticillium dahliae*. Our results show that Dim2, but not Dnmt5 or the putative sexual-cycle related DNA methyltransferase Rid, is responsible for nearly all DNA methylation. Single or double DNA methyltransferase mutants did not show altered development, virulence, or transcription of genes or transposons. In contrast, *Hp1* and *Dim5* mutants that are impacted in chromatin-associated processes upstream of DNA methylation are severely affected in development and virulence and display extensive transcriptional reprogramming in specific hypervariable genomic regions (so-called lineage-specific (LS) regions) that contain genes associated with host colonization. As these LS regions are largely devoid of DNA methylation and of Hp1- and Dim5-associated heterochromatin, the differential transcription is likely caused by pleiotropic effects rather than by differential DNA methylation. Overall, our study suggests that Dim2 is the main DNA methyltransferase in *V. dahliae* and, in conjunction with work on other fungi, is likely the main active DNMT in ascomycetes, irrespective of *Dnmt5* presence. We speculate that Dnmt5 acts under specific, presently enigmatic, conditions or, alternatively, acts in DNA-associated processes other than DNA methylation.

## INTRODUCTION

Transcriptional control is important for regulating developmental processes and environmental responses. In eukaryotes, transcriptional control is achieved through transcription factor-mediated and epigenetic mechanisms, the latter affecting DNA accessibility and altering interactions between DNA and various proteins (Burdge *et al*., 2007; Chen, 2007; Huang *et al*., 2018). Eukaryotic DNA associates with histone-protein complexes to form nucleosomes that are the main constituents of chromatin, a highly ordered DNA-structure (Luger *et al*., 1997). DNA accessibility for the transcriptional machinery is regulated in part by chemical modifications to histones that can alter chromatin structure or nucleosome positioning, and by direct DNA modifications that can alter transcription factor-binding sites (Slotkin & Martienssen, 2007). One such DNA modification is mediated by DNA methyltransferases (DNMT) that covalently add a methyl group to the 5^th^ carbon of a cytosine residue (5-methylcytosine, 5mC) (Bird, 1992). Cytosine methylation can occur in symmetric CG or CHG genomic contexts, or in the asymmetric CHH genomic context, where H stands for either A, C or T. In general, 5mC occurs more commonly at symmetric sites because maintenance methylation can cause methylation of daughter strands during DNA-replication, whereas asymmetric sites require *de novo* methylation (Law & Jacobsen, 2010). In mammals, DNA methylation is largely restricted to CG sites, while plants and fungi show methylation in each of the genomic contexts (Schmitz *et al*., 2019).

Compared to animal and plant genomes, fungi typically have smaller and less complex genomes, and they serve as important eukaryote models for various cellular processes including DNA methylation (Galagan *et al*., 2005). Much of the initial research on DNA methylation in fungi was performed in the saprophytic ascomycete fungus *Neurospora crassa*. In *N. crassa*, DNA methylation is restricted to transposable elements and is dependent on a single DNMT, Deficient In Methylation-2 (Dim2), an ortholog of Human Dnmt1 that performs *de novo* as well as maintenance methylation (Kouzminova & Selker, 2001). Dim2 operates in a complex with Heterochromatin Protein-1 (Hp1) that recognizes and directs DNA methylation to genomic regions marked by tri-methylation of histone 3 lysine 9 (H3K9me3) that is deposited by the histone methyltransferase Deficient In Methylation-5 (Dim5) (Tamaru & Selker, 2001; Freitag *et al*., 2004). Besides Dim2, *N. crassa* encodes another DNMT domain-containing protein of the fungal-specific class Dnmt4, named Repeat-Induced Point Mutation (RIP)-Defective (Rid), which is only active during sexual reproduction (Cambareri *et al*., 1989; Lewis *et al*., 2009). However, Rid has not been shown to methylate DNA, but is required for the RIP mechanism that can induce C to T mutations in duplicated genomic regions, including transposons (Cambareri *et al*., 1989; Lewis *et al*., 2009). Similar to *N. crassa*, the ascomycete plant pathogenic rice blast fungus *Magnaporthe oryzae* encodes orthologues of Dim2 and Rid. However, in contrast to *N. crassa* Rid, *M. oryzae* Rid displays DNA methylation activity, albeit with lower activity than Dim2 (Ikeda *et al*., 2013; Jeon *et al*., 2015). The opportunistic human pathogenic basidiomycete *Cryptococcus neoformans* encodes neither Dim2 nor Rid, but relies on an ortholog of Human Dnmt5 for DNA methylation (Huff & Zilberman, 2014). *C. neoformans* Dnmt5 can methylate DNA through direct binding to H3K9me3 or through association with the Hp1 homolog Swi6 (Catania *et al*., 2020). Additionally, *C. neoformans* Dnmt5 performs maintenance methylation through association with the 5mC-reader Uhrf1 that recognizes hemi-methylated CG sites (Catania *et al*., 2020). Recent phylogenetic analyses of DNMTs across the fungal kingdom revealed extensive diversity in the DNMT repertoires, with only few (less than 10%) species containing either both Dim2 and Rid, or only Dnmt5, whereas many contain the combination of Dim2, Rid and Dnmt5 (Bewick *et al*., 2019). Thus, our knowledge on DNA methylation in fungi has been primarily based on species that are not representative for the typical DNMT repertoire of most fungi.

*Verticillium dahliae* is a xylem-invading, soil-borne ascomycete fungus that causes Verticillium wilt disease on hundreds of plant species (Fradin & Thomma, 2006; Klosterman *et al*., 2009). Sexual reproduction has not been reported for *V. dahliae* that is presumed to mainly reproduce asexually (de Jonge *et al*., 2013). Recently, we demonstrated that DNA methylation in *V. dahliae* requires Hp1 and is restricted to H3K9me3-enriched transposons that localize mainly in evolutionary stable core genomic regions that are typically shared across different *V. dahliae* strains, including centromere regions (Cook *et al*., 2020; Seidl *et al*., 2020). In contrast to stable core regions, genomic regions that are important for adaptation show extensive presence-absence polymorphisms between *V. dahliae* strains, and are therefore designated as lineage-specific (LS) (Klosterman *et al*., 2011; de Jonge *et al*., 2013; Faino *et al*., 2016; Depotter *et al*., 2019; Cook *et al*., 2020). Many genes that play critical roles in host colonization reside in LS regions (Klosterman *et al*., 2011; de Jonge *et al*., 2012, 2013; Faino *et al*., 2016). LS regions are enriched in transposons that typically lack DNA methylation, which corresponds with increased transcriptional activity when compared with transposons in the core genome (Cook *et al*., 2020). Interestingly, the transcriptional activity of transposons seems instrumental for the evolution of LS regions (Faino *et al*., 2016), indicating that transposons in the core genome may carry DNA methylation to supress their transcriptional activity and to prevent genomic alterations that might reduce fitness. In this study, we investigated the contribution of various putative components of the methylation machinery on the physiology and biology of *V. dahliae* by performing bisulfite sequencing (BS-seq), transcriptomic analysis (RNA-seq), and functional studies on DNA methylation-associated genes.

## RESULTS

### The genome of *V. dahliae* encodes three putative DNA methyltransferases

Putative DNMTs in *V. dahliae* were identified using homology searches to known fungal DNMTs. We selected representative basidiomycete, ascomycete and phycomycete fungi that were previously shown to have DNA methylation, as well as ascomycete *Fusarium* species that are related to *Verticillium*. The predicted proteomes of the selected species were searched with a Hidden Markov Model (HMM) pfam model (PF00145) that is characteristic for Dnmt1, Dim2, Rid and Dnmt5. Whereas *N. crassa, M. oryzae* and *C. neoformans* possess either a combination of *Dim2* and *Rid*, or only *Dnmt5*, our analyses showed that several ascomycete species, including all ten species of the *Verticillium* genus, encode all three DNMTs (Figure 1, Figure S1). Thus, *Verticillium* spp. encode the most commonly shared DNMT complement as observed in ascomycete fungi (Bewick *et al*., 2019).

**Figure 1.**
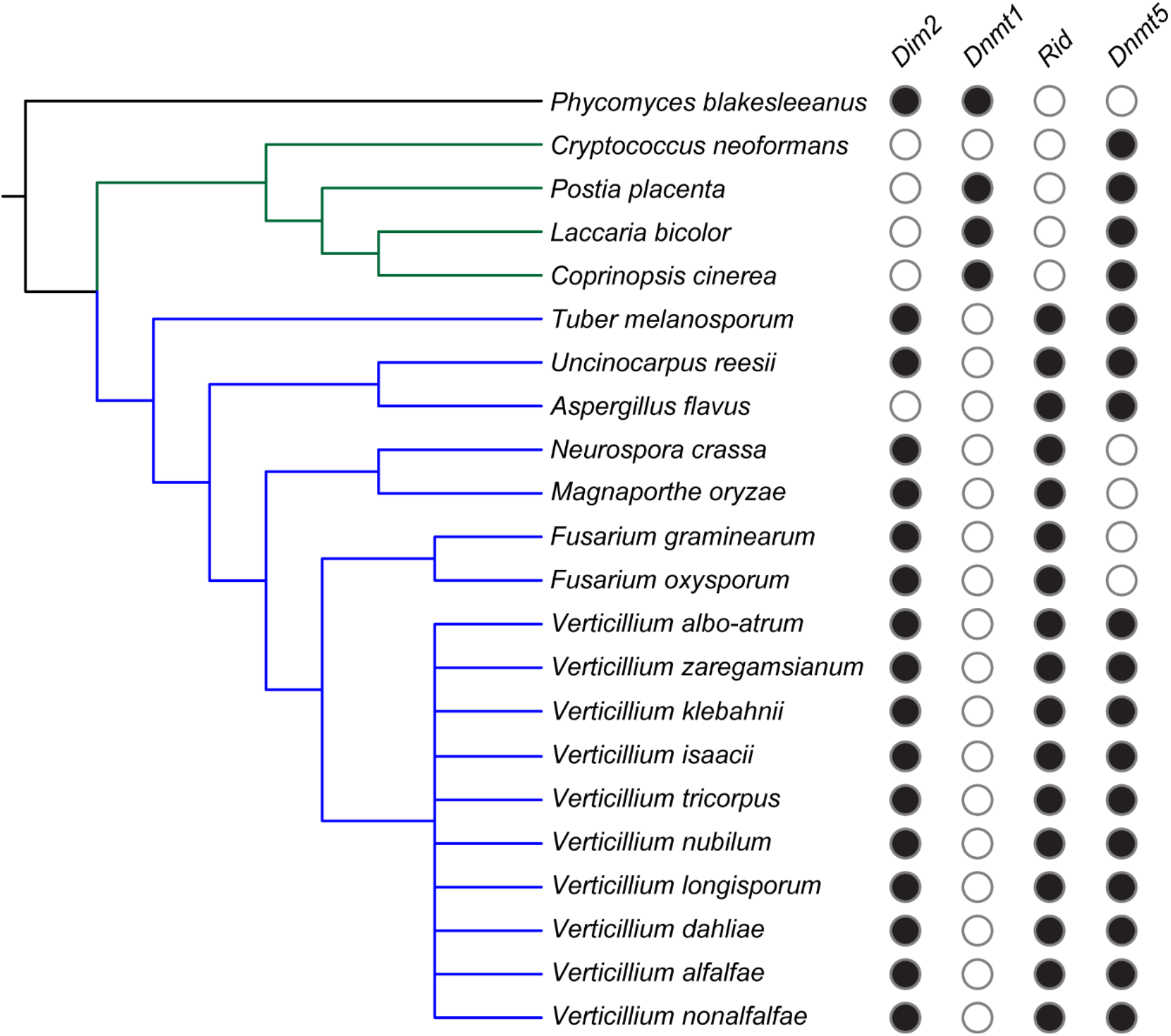
Presence of putative 5mC DNA methyltransferases in various fungi. Phylogenetic tree showing a phycomycete (black line), basidiomycetes (green lines) and ascomycetes (blue lines). Filled circles indicate presence of the corresponding DNA methyltransferase as indentified in Figure S1.

### DNA methyltransferase mutants are not affected in growth and virulence

As *V. dahliae* encodes three potential DNA methyltransferases, we sought to determine their activity and impact on development and virulence. To this end, we constructed deletion mutants for each DNMT gene, Δ*Dim2*, Δ*Dnmt5* and Δ*Rid*, as well as the Δ*Dim2*Δ*Dnmt5* double mutant in *V. dahliae* strain JR2. We did not generate double mutants with Δ*Rid* as well as a triple mutant because we anticipated, based on the lack of sexual reproduction in *V. dahliae* (de Jonge *et al*., 2013), that the role of *Rid* in DNA methylation, if any, would be negligible. We furthermore generated the H3K9 histone methyltransferase deletion mutant Δ*Dim5*, and used Δ*Hp1* (Cook *et al*., 2020), the DNA methyltransferase-complex member that recognizes H3K9me3 (Tamaru & Selker, 2001; Freitag *et al*., 2004).

Growth impacts were assessed for each strain under axenic growth as determined by colony size, spore production and morphology. Whereas all DNMT mutants displayed similar growth rates, spore production and colony morphology when compared with the wild-type strain, both Δ*Hp1* and Δ*Dim5* displayed decreased radial growth, with Δ*Dim5* being more strongly affected than Δ*Hp1* (Figure 2A, C). However, whereas Δ*Hp1* produced statistically significant fewer spores, Δ*Dim5* produced similar amounts of spores as wild-type *V. dahliae* when also considering their respective colony sizes (Figure 2B). This is likely due to Δ*Hp1* growing relatively flat, similar to wild-type *V. dahliae*, while Δ*Dim5* colonies display a severely crinkled surface, leading to an increased surface area on the same area of cultivation medium (Figure 2C). Both Δ*Hp1* and Δ*Dim5* displayed reduced pigmentation when compared with wild-type *V. dahliae* (Figure 2C)

**Figure 2.**
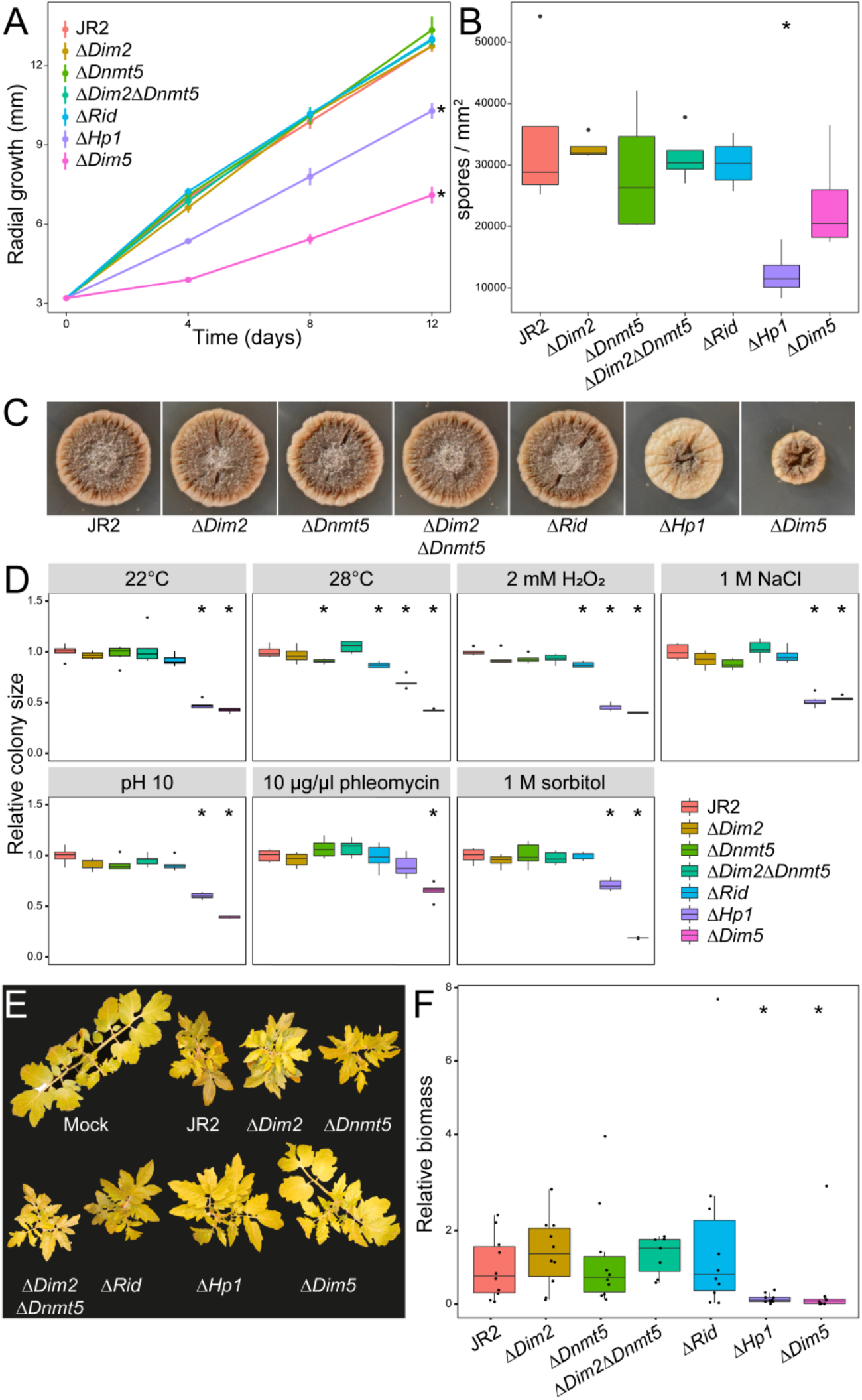
DNMT mutants of *Verticillium dahliae* do not show altered growth under axenic conditions, stress, or host colinization. A) Radial growth of wild-type and mutants over 12 days, with B) number of spores produced per mm^2^ of colony and C) pictures showing representative colony morphology after 12 days of growth. D) Colony area of wild-type and mutants subjected to various stress agents, relative to average colony area of wild-type. E) Representative pictures of infected tomato plants at 21 days after inoculation, with F) biomass of wild-type and mutants, relative to wild-type infection. Statistically significant differences from wild-type (Wilcoxon Signed Rank, p < 0.01) are indicated with asterisks. For (A), statistical tests were only performed on colony diameter at 12 dpi.

The deletion mutants were also assessed for growth under abiotic stress conditions by axenically culturing all the strains at elevated temperature, or in the presence of osmotic, oxidative and genotoxic stress agents. Under these conditions, the DNMT mutants grew similar as the wild-type strain, while both Δ*Hp1* and Δ*Dim5* displayed reduced growth (Figure 2D). Interestingly, however, Δ*Hp1* grew similar as the wild-type strain when exposed to the genotoxic compound phleomycin despite its growth retardation under all other conditions tested (Figure 2D).

The ability to infect tomato plants was also assessed for all mutants. Tomato plants inoculated with any of the DNMT mutants displayed severe stunting at a level similar to plants inoculated with the wild-type strain (Figure 2F). Fungal biomass measurements on the infected plants confirmed that fungal colonization by the DNMT mutants was similar to that of the wild-type strain (Figure 2E). In contrast, Δ*Hp1* and Δ*Dim5* displayed significantly reduced tomato infection, evidenced by a similar canopy area of plants inoculated with these mutants when compared with mock-inoculated plants, as well as by the finding that inoculated plants contained only low amounts of fungal biomass (Figure 2E). Arguably, the observation of significantly reduced plant infection for both Δ*Hp1* and Δ*Dim5* should be attributed to their compromised growth characteristics (Figure 2A).

### Dim2 is the main DNA methyltransferase in *V. dahliae*

To determine the role of the putative DNMTs in cytosine methylation in *V. dahliae*, whole-genome bisulfite sequencing was conducted on the wild-type strain, along with the *DNMT* and *Hp1* mutants. We recently reported that wild-type *V. dahliae* displays relatively low levels of DNA methylation, with an average of ∼0.4% methylation in CG and CHG context and essentially no DNA methylation in CHH context (Cook *et al*., 2020). DNA methylation in *V. dahliae* is restricted to particular inactive transposons that locate in condensed, H3K9me3-enriched, chromatin regions in the core genome, including those localized in centromeres (Figure 3A) (Cook *et al*., 2020; Seidl *et al*., 2020). We furthermore showed that the Δ*Hp1* mutant lost all DNA methylation, indicating that Hp1 is required for cytosine methylation and *V. dahliae* DNMTs cannot methylate DNA independently (Cook *et al*., 2020).

**Figure 3.**
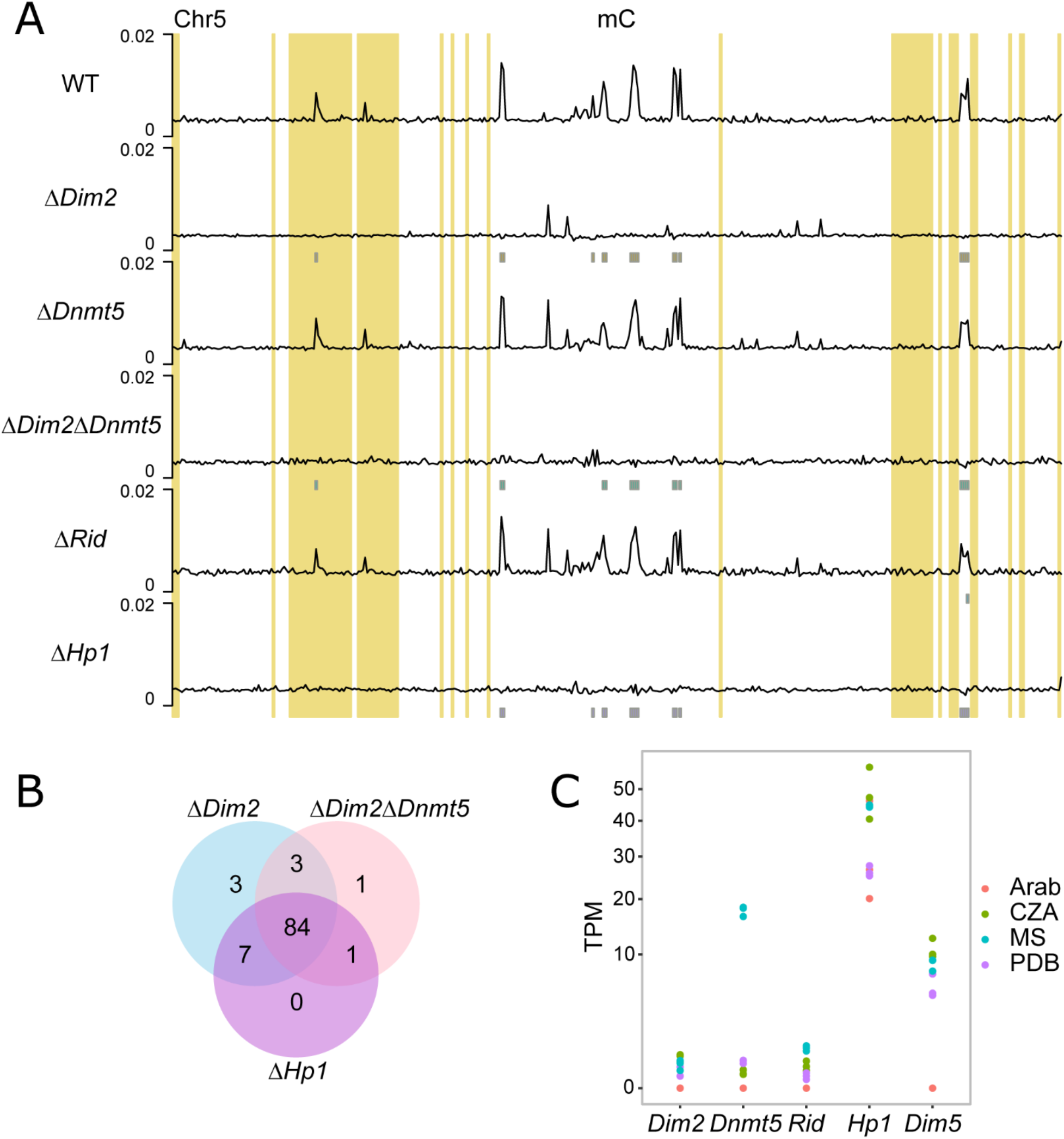
Dim2 is the main DNA methyltransferase in *V. dahliae*. A) Whole-chromosome plot displaying the fraction of methylated cytosines for non-overlapping 10 kb windows for wild-type, and *DNMT* and *Hp1* deletion mutants with chromosome 5 as an example. Grey boxes, displayed below the DNA methylation tracks, indicate the hypomethylated windows compared to the wild-type strain in CG and CHG context from Table 1. Previously defined LS regions (Cook *et al*., 2020) are highlighted in yellow. B) Overlap of hypomethylated windows in mutant strains showing severe loss of methylation. C) Expression (TPM values) of DNA methyltransferase genes *Dim2, Dnmt5* and *Rid*, as well as *Hp1* and *Dim5* of *V. dahliae* strain JR2 cultured in Czapec-Dox medium (CZA), half strength Murashige-Skoog medium (MS) and potato dextrose broth (PDB), and during Arabidopsis infection at 21 days post inoculation (Arab), in triplicates.

To study the extent to which the different *V. dahliae* mutants lost DNA methylation, we compared the bisulfite sequencing patterns over the genome in 10 kb windows and assessed the amount and location of hypomethylated windows when compared with the wild-type methylation pattern. Of the DNMT deletion mutants, Δ*Dim2* showed considerable loss of cytosine methylation, having 100 and 61 hypomethylated windows in the CG and CHG context, respectively (Table 1). As there is little methylation in CHH context, we combined the methylation data for CG and CHG context and also determined hypomethylation for the contexts simultaneously. This combination optimizes the number of potential methylated cytosines per window and therefore better captures differential methylation. In the combined contexts we observed 97 hypomethylated windows that locate at regions that have relatively high methylation percentages in wild-type *V. dahliae* (Table 1, Figure 3, Figure S5). Notably, additional regions showed reduced DNA methylation in Δ*Dim2*, yet these were not classified as hypomethylated because the methylation level was already low in the wild-type and therefore did not meet our criteria for calling hypomethylated regions (see methods for details). In contrast to the results for Δ*Dim2*, Δ*Dnmt5* and Δ*Rid* largely retained DNA methylation with only three and twelve windows being hypomethylated in CG context and fifteen and seventeen windows in CHG context, respectively (Table 1). When assessing CG and CHG methylation combined, Δ*Dnmt5* and Δ*Rid* have one and eight hypomethylated windows, respectively. Additionally, their genome-wide DNA methylation patterns are similar to the wild-type with no obvious loss of DNA methylation peaks (Figure 3A). The Δ*Dim2*Δ*Dnmt5* double mutant as well as Δ*Hp1* showed similar cytosine methylation levels over the genome as Δ*Dim2* and had hypomethylation of 93 and 99 windows in CG context and 59 and 65 windows in CHG context, respectively, and 89 and 92 hypomethylated windows when combining CG and CHG methylation data (Table 1, Figure 3A). Loss of methylation in the Δ*Dim2*, Δ*Dim2*Δ*Dnmt5* and Δ*Hp1* mutants largely occured in the same genomic regions, as 84 of the hypomethylated windows were shared between the mutants (Figure 3B). Even though the chromosome plots of the Δ*Dim2*Δ*Dnmt5* double mutant as well as Δ*Hp1* are similar to those of Δ*Dim2*, the few bins with slightly elevated methylation levels in Δ*Dim2* have decreased further (Figure 3A). This finding suggests that Dnmt5 has DNA methylation activity on particular genomic regions, albeit at a lower level. However, Δ*Dnmt5* does not display reduced methylation at the regions that remain slightly methylated in Δ*Dim2* (Figure 3A). Thus, if Dnmt5 has DNA methylation activity, it is redundant and secondary to the DNA methylation activity of Dim2. No windows were hypomethylated for CHH in any of the mutants (Table 1, Figure S4). The few bins with low levels of DNA methylation that locate in LS regions behave similar as those in core regions, in that they are hypomethylated in Δ*Dim2*, Δ*Hp1* and Δ*Dim2*Δ*Dnmt5* (Figure 3A). These results show that the methyltransferase Dim2 is responsible for the vast majority of detectable DNA methylation in *V. dahliae*.

**Table 1:** Number of 10 kb windows that are hypomethylated in *Verticillium dahliae DNMT* mutants relative to those in the wild-type strain.

| Genotype | Hypo-methylated windows (CG) | Hypo-methylated windows (CHG) | Hypo-methylated windows (CHH) | Hypo-methylated windows (CG and CHG) |
| --- | --- | --- | --- | --- |
| $\Delta Dim2$ | 100 | 61 | 0 | 97 |
| $\Delta Dnmt5$ | 3 | 15 | 0 | 1 |
| $\Delta Dim2\Delta Dnmt5$ | 93 | 59 | 0 | 89 |
| $\Delta Rid$ | 12 | 17 | 0 | 8 |
| $\Delta Hp1$ | 99 | 65 | 0 | 92 |

We compared *V. dahliae* Dnmt5 to the homolog in *C. neoformans*, where it is the sole active DNA methyltransferase (Catania *et al*., 2020). The two proteins share only 18% sequence similarity, but do share similar domain structures, except that the *V. dahliae* Dnmt5 lacks the N-terminal chromo-shadow domain found in *C. neoformans* Dnmt5 (Figure S6). This domain is responsible for the direct binding to H3K9me3, and this histone mark is required for DNA methylation, which could explain why we observed little Dnmt5 contribution to DNA methylation in *V. dahliae*. However, *C. neoformans* Dnmt5 can also bind H3K9me3 indirectly through Hp1 (Catania *et al*., 2020), and it is not clear if this is also the case for *V. dahliae*. The lack of DNA methylation by Dnmt5 cannot be explained by transcriptional activity, as *Dnmt5* is expressed higher than *Dim2* during cultivation in PDB (Figure 3B).

### Loss of DNA methylation does not affect transcriptional regulation

While the *Dim2* mutant loses nearly all DNA methylation (Table 1, Figure 3A), it displays wild-type-like growth *in vitro*, under stress conditions as well as during infection (Figure 2), suggesting that DNA methylation is not essential under these conditions. However, given that DNA methylation is mainly restricted to transposons (Cook *et al*., 2020), we anticipated that loss of DNA methylation could result in activated transcription at transposons. To address this, we performed RNA sequencing on axenically grown cultures of all DNMT mutants, as well as the *Hp1* and *Dim5* mutants. Consistent with the lack of DNA methylation at coding regions, all three single DNMT mutants and the double mutant showed differential expression of only few genes (< 10) (Figure 4A). Unanticipatedly, the four DNMT mutant strains similarly showed differential expression of only a few transposons (< 10) (Figure 4B). In contrast, the Δ*Hp1* and Δ*Dim5* mutant strains showed considerable differential expression of genes and transposons (Figure 4A). In total, 1,759 genes were induced and 688 repressed in Δ*Hp1*, and 1,638 genes were induced and 787 are repressed in Δ*Dim5* when compared with wild-type (Figure 4A). Furthermore, 320 transposons were induced and 29 were repressed in Δ*Hp1*, whereas 264 transposons were induced and 50 were repressed in Δ*Dim5* when compared with wild-type (Figure 4B). Analysis of the induced genes and transposons revealed a large overlap between the mutants, with 1,280 out of 1,638 (∼78%) of the induced genes and 191 out of 264 (∼72%) of the induced transposons shared between the two mutants (Figure 4A, B). This overlap is likely related to the functional link between these heterochromatin components, as Hp1 directs DNA methylation to H3K9me3 deposited by Dim5 (Freitag *et al*., 2004). To study whether the genes activated in Δ*Hp1* and Δ*Dim5* represent specific biological functions, we performed GO enrichment analysis on the 1,280 induced genes. Interestingly, genes encoding secreted proteins and proteins involved in transport and metabolic processes were overrepresented among the induced genes (Figure 4C), suggesting that the mutants impact expression of genes with roles in responses to the environment.

**Figure 4.**
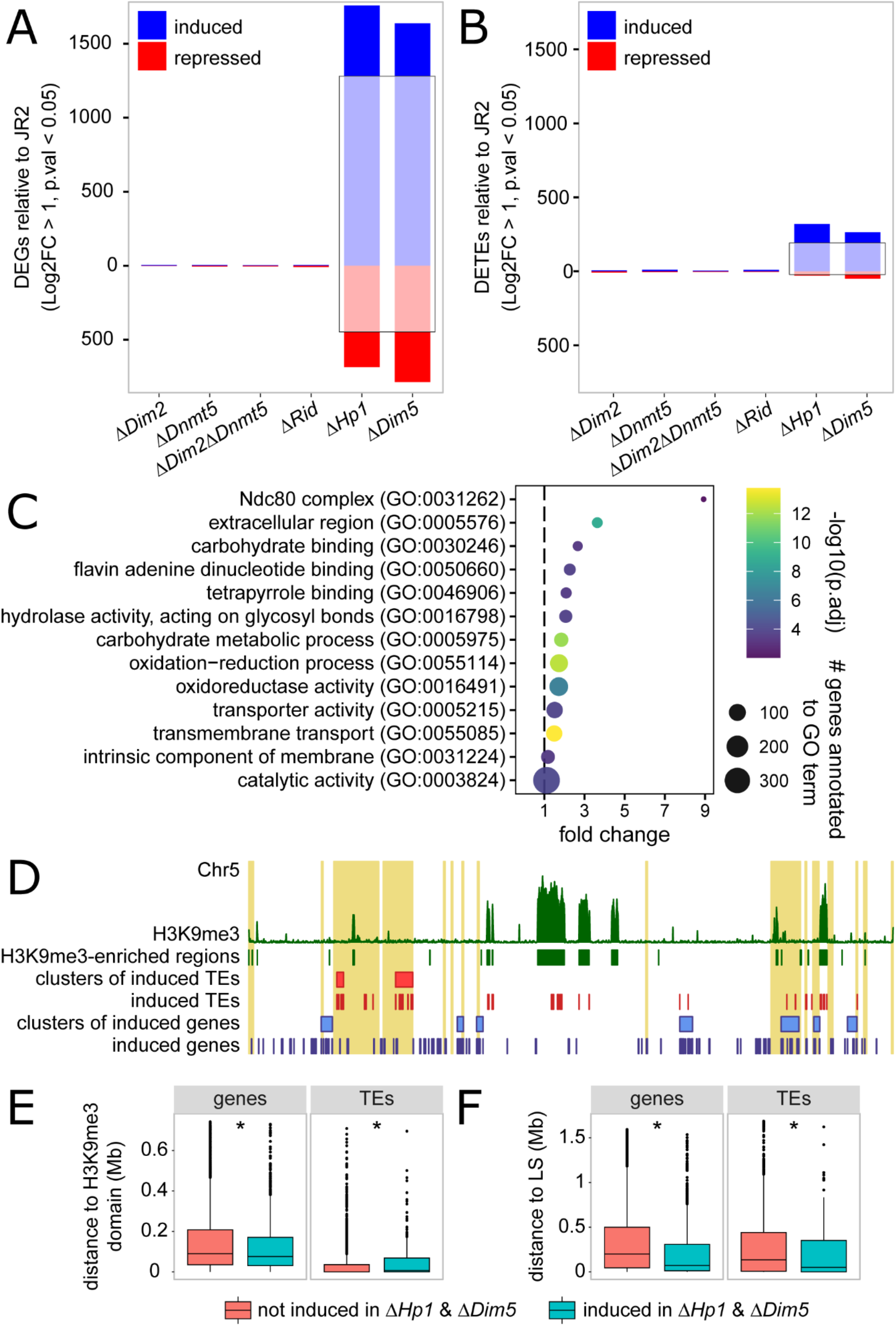
Genes and transposons that are induced in the *Verticillium dahliae Hp1* and *Dim5* mutants do not associate with H3K9me3-marked chromatin. Differentially expressed genes (A) and transposons (B) in the mutants relative to wild-type. Induced genes and transposons are indicated in blue, repressed genes and transposons in red. The number of genes and transposons that are induced and repressed in both the *Hp1* and *Dim5* mutants are indicated by opaque coloring in black rectangles. The amounts of genes (C) and transposons (D) that are induced in both the *Hp1* and *Dim5* mutants, and occur clustered in the genome (blue line) compared to a distribution of 1000 random equally sized sets of genes and transposons. (E) Whole-chromosome plot displaying the location of induced genes (in blue) and transposons (in red) on chromosome 5 as an example. Clusters of induced genes and transposons are indicated as blue and red rectangles, respectively. H3K9me3-ChIP signal along the chromosome is indicated in green in the upper track. LS regions (Cook *et al*., 2020) are highlighted in yellow. The minimal distance of genes and transposons to H3K9me3-enriched genomic regions (F) and to LS regions (G). Asterisks indicate statistical differences (Wilcoxon signed rank test, p<0.01) between genes and TEs induced in both the Δ*Hp1* and Δ*Dim5* mutants and those that are not induced in the mutants.

As Hp1 functions downstream of Dim5 during DNA methylation, we expect that the induction of genes as well as transposons in both mutants may be due to reduced recruitment of repressive complexes to previously silenced chromatin regions because either H3K9me3 or Hp1 is lacking. Based on H3K9me3 ChIP-seq, we found that approximately 2.1 Mb (∼6%) of the genome is associated with H3K9me3 that occurs in 621 enriched genomic regions, of which 38 are larger than 10 kb (Figure S7). To study whether the induced genes and transposons localize in these H3K9me3 domains, we investigated the occurrence of physical clustering of the 1,280 induced genes and the 191 induced transposons. Our hypothesis was that Dim5 deposited H3K9me3 and associated Hp1 mediate transcriptional silencing in physical proximity to H3K9me3 domains. As such, we expected that genes and transposons induced in Δ*Hp1* and Δ*Dim5* occured in clusters and that these clusters would be in proximity to H3K9me3-enriched genomic regions. We identified 57 clusters containing 559 of the 1,280 (∼44%) induced genes and five clusters containing 49 of the 191 (∼26%) induced transposons, which is more than expected by chance, as measured from 1,000 random sets of 1,280 genes (p<0.001) (Figure S8) and 49 transposons (p = 0.024) (Figure S9). H3K9me3 domains contain numerous transposons (1,034 out of 2,574, ∼40%) and only few genes (76 out of 11426, ∼0.6%) (Figure 4D, Figure S10) (Cook *et al*., 2020). Next, we calculated the distance of each gene and transposon to the closest H3K9me3 domain and associated this to induced and non-induced genes and transposons. When considering the smallest distance to H3K9me3 domains, we observed that induced genes are slightly closer to H3K9me3 domains than non-induced genes (Figure 4E). In contrast, induced transposons are slightly further from H3K9me3 domains than non-induced transposons (Figure 4E). Additionally, when considering presence of transposons in H3K9me3 domains, we observed that significantly fewer induced transposons, 53 out 191 (27.7%), than non-induced transposons, 981 out of 2,383 (41.2%), locate in H3K9me3 domains (Fisher’s exact test, p = 0.0126). The relatively minor enrichment of induced genes near H3K9me3 domains and the enrichment of induced transposons away from H3K9me3 domains suggests that the transcriptional changes in Δ*Hp1* and Δ*Dim5* are not due to reduced H3K9me3 and Hp1 association. As we demonstrated that the induced genes and transposons occur clustered in the genome (Figure S8, S9), we asked whether the clusters localize in specific genomic regions. As we found that induced genes are enriched for genes encoding secreted proteins and proteins involved in metabolic processes (Figure 4C), we speculated that the induced genes may be involved in processes related to plant infection. Such genes are typically located in LS regions of *V. dahliae*, which are enriched in transposons but are not associated with H3K9me3 (de Jonge *et al*., 2013; Cook *et al*., 2020). Therefore, we also tested whether the induced genes and transposons locate in proximity to LS regions. Intriguingly, genes and transposons induced in Δ*Hp1* and Δ*Dim5* are significantly closer to LS regions than non-induced genes and transposons (Figure 4D,F). Additionally, 197 out of 1,280 (15.4%) and 70 out of 191 (36.6%) of the induced genes and transposons locate in LS regions, which is significantly more than the 801 out of 10,146 (7.9%) and 463 out of 2,383 (19.4%) of non-induced genes (Fisher’s exact test, p <0.00001) and transposons (Fisher’s exact test, p <0.00001). Consequently, both genes and transposons that are induced in Δ*Hp1* and Δ*Dim5* reside significantly closer to LS regions than non-induced genes and transposons (Figure 4F). Since LS regions are not associated with H3K9me3, these findings suggest that the observed transcriptional changes are not directly related to loss of Hp1 binding at H3K9me3 domains, but rather through pleiotropic effects affecting transcription throughout the genome, and especially at LS regions.

## DISCUSSION

DNA methylation is essential for proper functioning of nuclear processes in many organisms (Bird, 1992), but various fungal species have lost or degraded their machinery for DNA methylation (Bewick *et al*., 2019). The most commonly found combination of DNMTs in ascomycete genomes is the presence of *Dim2, Rid* and *Dnmt5* (Bewick *et al*., 2019). As *Dim2* is the main DNA methyltransferase gene in fungal species that lack *Dnmt5*, and vice versa, it is relevant to study the importance of these DNA methyltransferase genes in fungal species that carry both. The fungal pathogen *Z. tritici* carries all three DNMTs and, similar to *V. dahliae*, loss of *Dim2* almost completely abolishes DNA methylation (Möller *et al*., 2020), indicating that Dnmt5 and Rid have little to no DNA methylation activity. However, a low residual DNA methylation signal remains in *Dim2* mutants of *V. dahliae* and *Z. tritici*, which may be due to a low degree of Dnmt5 activity (Figure 3; Möller *et al*., 2020). Our results indicate that *Dnmt5* is more highly expressed during growth in nutrient-limited media, a type of environmental stress. It is possible *Dnmt5* may be more active and cause differential DNA methylation during specific growth conditions not tested here, an occurrence that has been observed for DNMTs in several plant and animal species (Meaney & Szyf, 2005; Dowen *et al*., 2012). Differential expression of DNMTs has previously been observed in the entomopathogenic ascomycete fungus *Cordyceps militaris* that contains orthologues of *Dim2* and *Rid* (Wang *et al*., 2015). Whether *V. dahliae* Dnmt5 plays such a role requires further study.

The importance of DNA methylation in fungi that are able to perform DNA methylation remains unclear. Deletion of functional components of DNA methylation did not result in clear phenotypic alterations in *N. crassa* or the necrotrophic plant pathogenic fungus *Botrytis cinerea*, while deletion of *Dim2* in *M. oryzae* leads to aberrant colony morphology and compromised conidiospore formation (Kouzminova & Selker, 2001; Jeon *et al*., 2015; Zhang *et al*., 2016). In *Z. tritici*, strains collected in the centre of origin of its wheat host carry DNA methylation, while strains collected in Europe contain mutated *Dim2* copies that lack DNA methylation (Möller *et al*., 2020). The *Z. tritici* strains that lack a functional copy of *Dim2* are at least as virulent as strains that perform DNA methylation (Haueisen *et al*., 2019), suggesting that the recent loss of DNA methylation in these *Z. tritici* strains does not negatively affect their infection biology. Our study reveals that DNA methylation in *V. dahliae* is not essential for growth and infection. Moreover, we show that loss of DNA methylation does not result in altered expression of genes or transposons, an observation that could be explained by DNA methylation co-localizing with H3K9me3, which is likely sufficient for heterochromatin formation and transcriptional silencing in the absence of DNA methylation. This is further supported as the H3K9me3-deficient *V. dahliae Dim5* mutant showed significant differential expression of genes and transposons. Interestingly, some genes and transposons induced in this mutant were located in LS regions that are not labelled with H3K9me3 in wild-type *V. dahliae*, suggesting that the removal of H3K9me3 leads to pleiotropic effects in unrelated genomic regions. Similar effects on gene and transposon expression occurs in the *V. dahliae Hp1* mutant, indicating that the differential expression observed in the *Hp1* and *Dim5* mutants are related to disrupted Hp1 functioning. In *N. crassa*, and also fission yeast *Schizosaccharomyces pombe* that lacks DNA methylation, Hp1 was found to be involved in the formation of H3K9me3-associated heterochromatin (Freitag *et al*., 2004; Hayashi *et al*., 2012; Gessaman & Selker, 2017). As such, it is possible that the transcriptional changes in the *V. dahliae Hp1* and *Dim5* mutants are due to pleiotropic effects of chromatin de-condensation.

Considering that experimental and natural loss of DNA methylation in various fungi does not seem to affect their proliferation, it is remarkable that the vast majority of fungal species have retained DNA methylation. One explanation for the role of DNA methylation in fungi, which accounts for the lack of reported phenotypes, is that it serves in maintaining genome integrity during evolution. In this way, DNA methylation does not functionally regulate transcription *per se*, but works in conjunction with H3K9me3 to minimize the impact of TEs in the genome. One possible mechanism is that DNA methylation may have persistent effects on transposon activity through spontaneous deamination of methylated cytosines, resulting in C to T mutations (Holliday & Grigg, 1993). The deamination process is considered an important driver of mutations in *Z. tritici* transposons as recently shown in an experimental evolution experiment in which a DNA methylation competent strain had increased in C to T mutations compared to the strain lacking DNA methylation (Möller *et al*., 2020). Interestingly, in *V. dahliae* we previously observed that transposons that carry DNA methylation contain more C to T mutations than unmethylated transposons (Cook *et al*., 2020). Typically, such C to T mutations are also caused by the RIP mechanism, which relies on the Rid DNA methyltransferase that is active during sexual cycles in *N. crassa* (Cambareri *et al*., 1989; Lewis *et al*., 2009). However, since *V. dahliae* is presumed to reproduce asexually, it may be more likely that C to T mutations in transposons are caused by spontaneous deamination. These results support that the main role for DNA methylation in fungi might be to aid in transposon sequence degradation over time, not to directly supress transcriptional activity. Alternatively, it is possible DNA methylation is important for inhibiting transcriptional activity of transposons during specific developmental or cell-cycle stages which have not been reported or observed to date.

Our results show that although *V. dahliae* encodes multiple DNMTs, only *Dim2* seems to be essential for DNA methylation. As *Dim2, Dnmt5* and *Rid* are wide-spread among ascomycetes, it is likely that their combined presence is an ancestral state (Bewick *et al*., 2019). Even though only four ascomycete species have been studied with respect to the contribution of their DNMTs to DNA methylation so far, these studies suggest that species, irrespective of the presence or absence of *Dnmt5*, utilize Dim2 as the main DNMT (this study; Kouzminova & Selker, 2001; Jeon *et al*., 2015; Möller *et al*., 2020). Additional research is needed to determine if Dnmt5 plays a role in DNA methylation, or possibly another pathway in these species, or if its presence is the remnant of an ancestral state that is not strongly selected against.

## MATERIALS AND METHODS

### Assessment of DNMT occurrence

To assess the presence of DNA methyltransferases in a selection of fungal species with confirmed DNA methylation performance, we downloaded predicted proteomes of *Aspergillus flavus* strain NRRL_3357 (AFL2T), *Coprinopsis cinerea* strain Okayama-7#130 (CC1G), *Cryptococcus neoformans* strain H99 (CNAG), *Fusarium graminearum* strain PH-1 (FGSG), *F. oxysporum* strain 4287 (FOXG), *Laccaria bicolor* strain S238N-H82 (lacbi2), *Magnaporthe oryzae* strain MG8 (MGG), *Neurospora crassa* strain OR74a (NCU), *Phycomyces blakesleeanus* strain NRRL 1555(-) (Phybl2), *Postia placenta* strain MAD698 (pospl1), *Tuber melanosporum* strain Mel28 (Tubme1), *Uncinocarpus reesii* strain 1704 (URET) and the *Verticillium albo-atrum* PD747, *V. alfalfae* PD683, *V. dahliae* JR2, *V. isaacii* PD618, *V. klebahni* PD401, *V. longisporum* PD589, *V. nonalfalfae* T2, *V. nubilum* 397, *V. tricorpus* PD593 and *V. zaregansianum* PD739. The predicted proteomes were scanned for the presence of a DNA methyltransferase domain with hmmsearch with --cut_ga option. Identified proteins were visually inspected. To check for presence of additional not-annotated DNA methyltransferase homologs of *Dim-2, Dnmt5* and *Rid* that were initially not annotated in the predicted proteomes, we manually assessed the genomes using TBLASTN and Augustus. Phylogenetic trees were constructed using IQ-tree.

### Fungal growth and mutant generation

*V. dahliae* strain JR2 (CBS 143773; Faino et al., 2015) was maintained on potato dextrose agar (PDA) (Oxoid, Thermo Scientific, CM0139) and grown at 22°C in the dark. The Δ*Dim2*, Δ*Dnmt5*, Δ*Rid* and Δ*Dim5* single deletion mutants and the Δ*Dim2*-Δ*Dnmt5* double mutant were constructed as previously described (Santhanam, 2012). Briefly, for all genes except *Dnmt5*, genomic DNA regions flanking the 5’ and 3’ ends of the coding sequences were amplified with PCR using primers listed in Table S1 and cloned in to the pRF-HU2 vector (Frandsen *et al*., 2008), using USER enzyme following the manufacturer’s protocol (New England Biolabs, MA, USA). For *Dnmt5*, the 5’ and 3’ amplicons were cloned into vector pRF-NU2, a custom-made pRF-HU2 variant, containing the NAT-cassette for selection on nourseothrycin. Sequence-verified vectors were transformed into *Agrobacterium tumefaciens* strain AGL1 used for *V. dahliae* conidiospore transformation as described previously (Santhanam, 2012). *V. dahliae* transformants that appeared on hygromycin B or nourseothrycin (for *Dnmt5*) were transferred to fresh PDA supplemented with hygromycin B or nourseothrycin after five days. Putative transformants were screened using PCR to verify deletion of the target gene sequence (Table S2) when compared with positive amplification from the wild-type strain. To further confirm integration of the selectable marker at the locus of interest, another round of PCR was conducted in which one primer was position adjacent to the deleted genomic region, and the other primer was designed to bind a portion of the inserted vector DNA (Table S3). In this manner, deletion mutants were confirmed to lack the gene of interest and contain the selectable marker at the locus of interest. Generation of the *Hp1* deletion mutant was conducted in the same way and described previously (Cook *et al*., 2020).

### Growth and inoculation assays

To check for aberrant growth phenotypes of the generated mutants, all strains were cultured as described above. To this end, conidiospores were harvested in sterile water and brought to a final concentration of 10^6^ conidiospores per mL. Subsequently, 10 µL of conidiospore suspension, containing 10^4^ conidiospores, was deposited in the middle of a 90 mm Petri dish containing 20 ml of PDA. Plates were stored at 22°C in the dark and colony diameter was measured in perpendicular directions after 4, 8 and 12 days of growth. After twelve days of growth all newly formed conidiospores were harvested in 1 mL of water and counted using a hemocytometer.

Stress assays were performed by spotting 5 µL conidiospore suspension containing 5×10^3^ conidiospores on PDA without supplement, or on PDA supplemented with 1 M NaCl, 1 M Sorbitol, 2 mM H_2_O_2_ or 10 µg/µL phleomycin, and on PDA adjusted to pH 10. Plates were incubated at 22°C in the dark, apart from one set of PDA plates without supplement that was incubated at 28°C to assess heat stress responses. Pictures were taken after 6 or 10 days, depending on wild-type colony development, and colony size was determined using ImageJ software with custom settings for each stress condition.

Infection assays were performed using root dip inoculation in a conidiospore suspension of 10^6^ spores per mL on 10-day-old seedlings of tomato cultivar Moneymaker. Stems of infected plants were harvested at 21 days after inoculation, cut in small pieces, frozen in liquid nitrogen and ground by reciprocal shaking in a MixerMill MM 400 (Retsch, Haan, Germany). DNA was isolated incubating the ground powder with 800 µL of CTAB lysis buffer at 65°C for 1 hour, followed by addition of 400 µL chloroform/IAA (24:1), vigorous shaking and centrifuging for 5 minutes at ∼13,000 RCF. DNA was precipitated from the aqueous layer with isopropanol and the precipitate was washed with 70% ethanol. The fungal biomass in the stem tissue was determined with real-time PCR using *V. dahliae* ITS-specific and tomato GAPDH-specific primer sets (Table S4).

### Bisulfite sequencing and analysis

The *V. dahliae* wild-type strain, Δ*Dim2*, Δ*Dnmt5*, Δ*Dim2* / Δ*Dnmt5*, Δ*Rid* and Δ*Hp1* were grown in potato dextrose broth (PDB) for three days, strained through miracloth (22 μm) (EMD Millipore, Darmstadt, Germany), pressed to remove excess liquid, flash frozen in liquid nitrogen and ground to powder with a mortar and pestle. Genomic DNA was isolated as described above and sent to the Beijing Genome Institute (BGI, Hong Kong, China) for bisulfite conversion, library construction and Illumina sequencing. Briefly, the DNA was sonicated to a fragment range of 100-300 bp, end-repaired and methylated sequencing adapters were ligated to 3’ ends. The EZ DNA Methylation-Gold kit (Zymo Research, CA, USA) was employed according to manufacturer’s guidelines for bisulfite conversion of non-methylated DNA. Libraries were paired-end 100 bp sequenced on an Illumina HiSeq 2000 machine.

Whole-genome bisulfite sequencing reads were analyzed using the BSMAP pipeline (v. 2.73) and methratio script (Xi & Li, 2009). The results were partitioned into CG, CHG and CHH cytosine sites for analysis. Only cytosine positions containing more than 4 sequencing reads were included for analysis. BSMAP datasets were further analyzed using MethylKit (v. 1.12.0) (Akalin *et al*., 2012). Methylation levels were summarized as the number of methylated cytosines divided by the total number of sequenced cytosines per 10 kb window. Hypomethylated windows in the mutants were determined by comparing corresponding 10 kb windows between mutants and wild-type and selecting windows with meth.diff value < -1 and a qvalue < 0.01. Genome plots displaying methylation data were generated using karyoploteR (v. 1.12.4) (Gel & Serra, 2017).

### Dnmt5 analysis and expression of DNA methyltransferase genes

The Dnmt5 amino-acid sequences of *V. dahliae* strain JR2 (VDAG_JR2_Chr1g14260) and *C. neoformans* var *grubii* strain H99 (CNAG_07552) were retrieved from EnsemblFungi, aligned using Uniprot.org/align and their domain structure was predicted using Interproscan. To assess expression of *Dim2, Dnmt5, Rid, Hp1* and *Dim5* in *V. dahliae* cultured in different growth media, wild-type *V. dahliae* was cultured in Czapec-Dox medium (CZA), half-strength Murashige-Skoog medium with vitamins and supplemented with 3% sucrose (MS), and potato dextrose broth (PDB) at 22°C at 160 RPM in the dark for six days and mycelium was collected for three replicates per growth medium and ground as described above. To obtain RNA-seq data from *V. dahliae* grown in planta, three-week-old *A. thaliana* (Col-0) plants were root dip inoculated in a conidiospore suspension of 10^6^ spores per mL for 10 minutes. After root inoculation, plants were grown in individual pots in a greenhouse for 21 days, under a cycle of 16 h of light and 8 h of darkness, with temperatures maintained between 20 and 22°C during the day and a minimum of 15°C overnight. Three pooled samples (10 plants per sample) of complete flowering stems were used for total RNA extraction, respectively. Total RNA from *in vitro* cultured mycelium and was isolated using Trizol (Thermo Fisher Science, Waltham, MA, USA) following the manufacturer’s guidelines. Following RNA re-suspension, contaminating DNA was removed using the TURBO DNA-free kit (Ambion, Thermo Fisher Science, Waltham, MA, USA) and RNA integrity was determined by separating 2 μL of each sample on a 2% agarose gel and quantified using a Nanodrop (Thermo Fisher Science, Waltham, MA, USA) and stored at -80°C until further use. Library preparation was carried out at BGI (BGI, Hong Kong, China) and 50 bp fragments were sequenced using the BGISEQ-500 platform. Sequenced reads were mapped to *V. dahliae* strain JR2 gene annotation using Kallisto quant (settings: --single -l 50 -s 0.001 --pseudobam) to obtain normalized TPM values (Bray *et al*., 2016).

### Transcriptional analysis of mutants in DNA methylation-associated genes

The *V. dahliae* wild-type strain JR2 (Faino *et al*., 2015), Δ*Dim2*, Δ*Dnmt5*, Δ*Dim2*-Δ*Dnmt5*, Δ*Rid*, Δ*Hp1* and Δ*Dim5* were cultured in PDB at 22°C at 160 RPM in the dark for six days. Mycelium collection, RNA-isolation and sequencing are performed for three replicates per growth condition as described above.

Differential gene and transposon expression between mutants and wild-type was determined by mapping sequencing reads to the *V. dahliae* strain JR2 gene and transposon annotation using TEtranscripts (Faino *et al*., 2015; Jin *et al*., 2015; Cook *et al*., 2020). Genes and transposons were considered differentially expressed when they displayed log2FoldChange of < -1 and a pvalue of < 0.05. Clustering of induced genes and transposons was determined using CROC (settings, for genes: -w 50000 -o 10000 -m 5, for transposons: -w 100000 -o 20000 -m 5) (Pignatelli *et al*., 2009). To analyze whether induced genes or transposons display more clustering than in random gene or transposon sets, 1,000 random selections of 1,280 genes and of 191 transposons were generated. These random gene and transposon sets were similarly analyzed using CROC. Overlap with H3K9me3 domains and LS regions (Cook *et al*., 2020) was assessed using bedtools closest (settings -d) (Quinlan & Hall, 2010).

### Data access

RNAseq data of Arabidopsis infection were submitted to the SRA database under the accession number: SRP149060.

## Acknowledgements

This work was supported by a PhD grant of the Research Council Earth and Life Sciences (project 831.15.002) to HMK and by Human Frontier Science Program Postdoctoral Fellowship (HFSP, LT000627/2014-L) to DEC. Work in the laboratories of M.F.S and B.P.H.J.T. is supported by the Research Council Earth and Life Sciences (ALW) of the Netherlands Organization of Scientific Research (NWO) and B.P.H.J.T is supported by the Deutsche Forschungsgemeinschaft (DFG, German Research Foundation) under Germany’s Excellence Strategy – EXC 2048/1 – Project ID: 390686111.

