## Supplemental Figures for "DNA methylation in *Verticillium dahliae* requires only one of three putative DNA methyltransferases, yet is dispensable for growth, development and virulence"

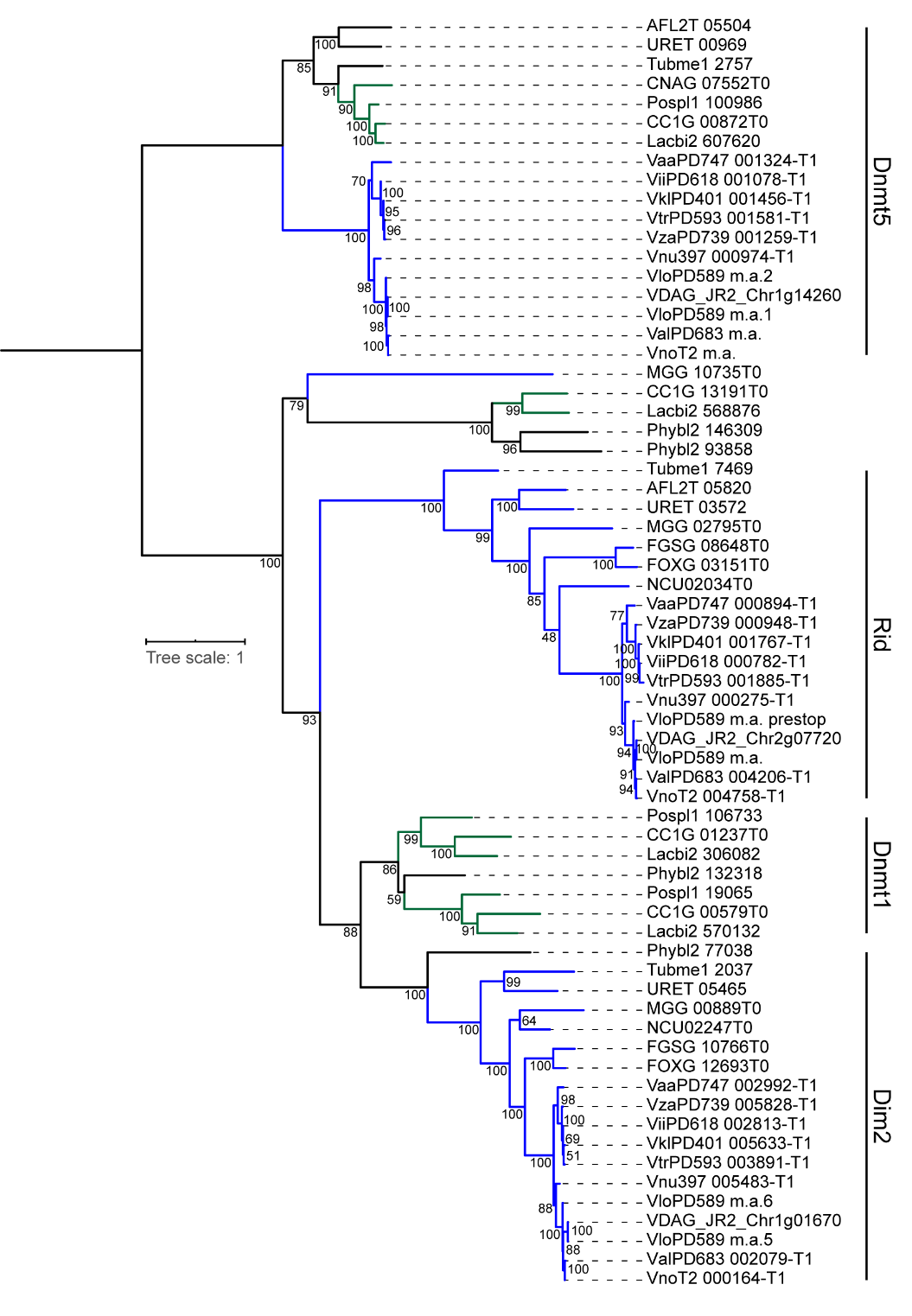

**Figure S1. Phylogenetic tree of DNA methyltransferases.** Gene codes containing m.a. were manually added, as they were missed in the predicted proteomes.

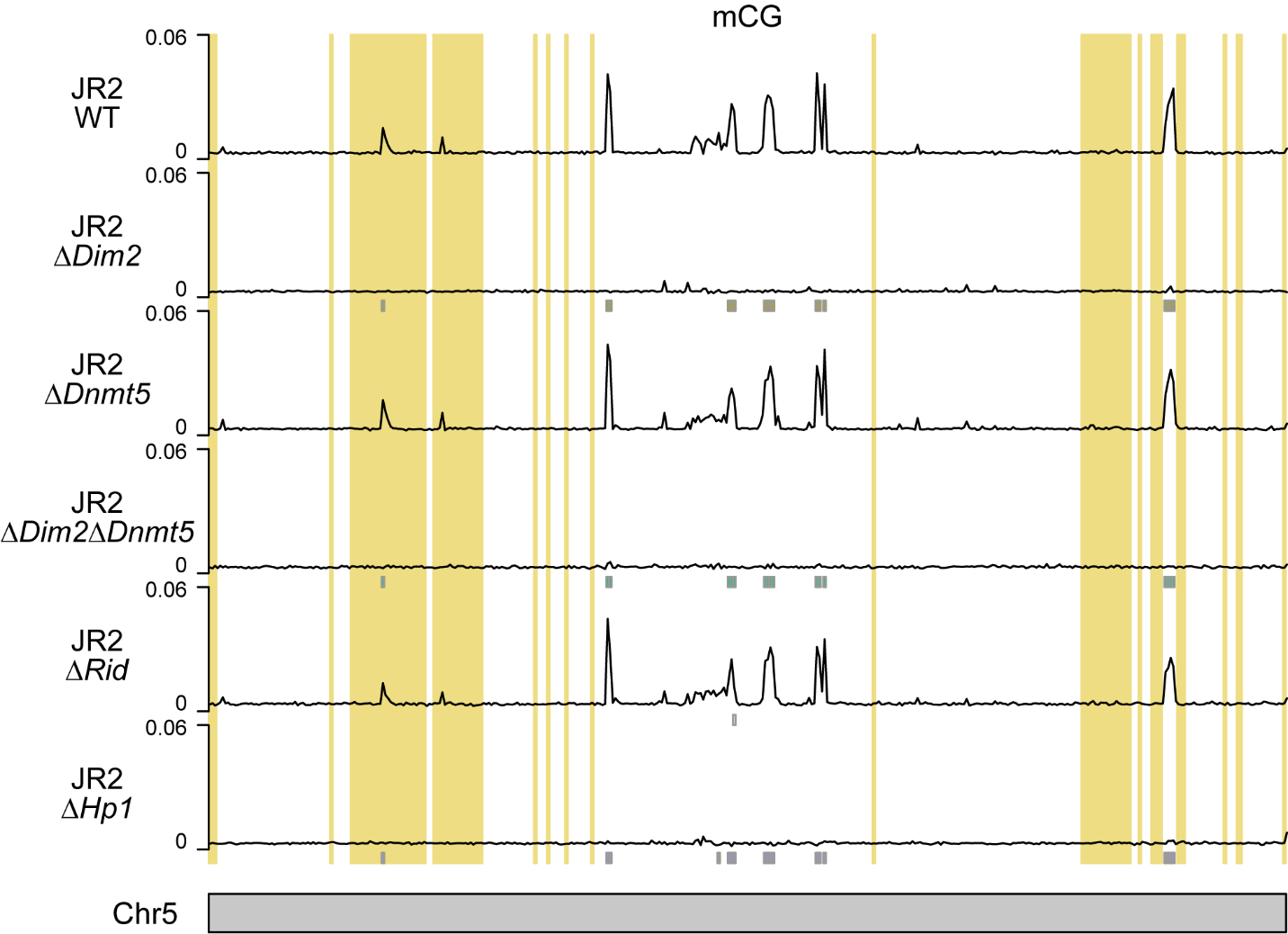

**Figure S2**. **DNA methylation in CG context.** Whole-chromosome plot displaying the fraction of methylated cytosines for non-overlapping 10 kb windows in CG context for WT, and DNA methyltransferase and Hp1 deletion mutants with chromosome 5 as an example. Grey boxes, displayed below the DNA methylation tracks, indicate the hypomethylated windows in CG context from Table 1. Previously defined LS regions (Cook et al., 2020) are highlighted in yellow.

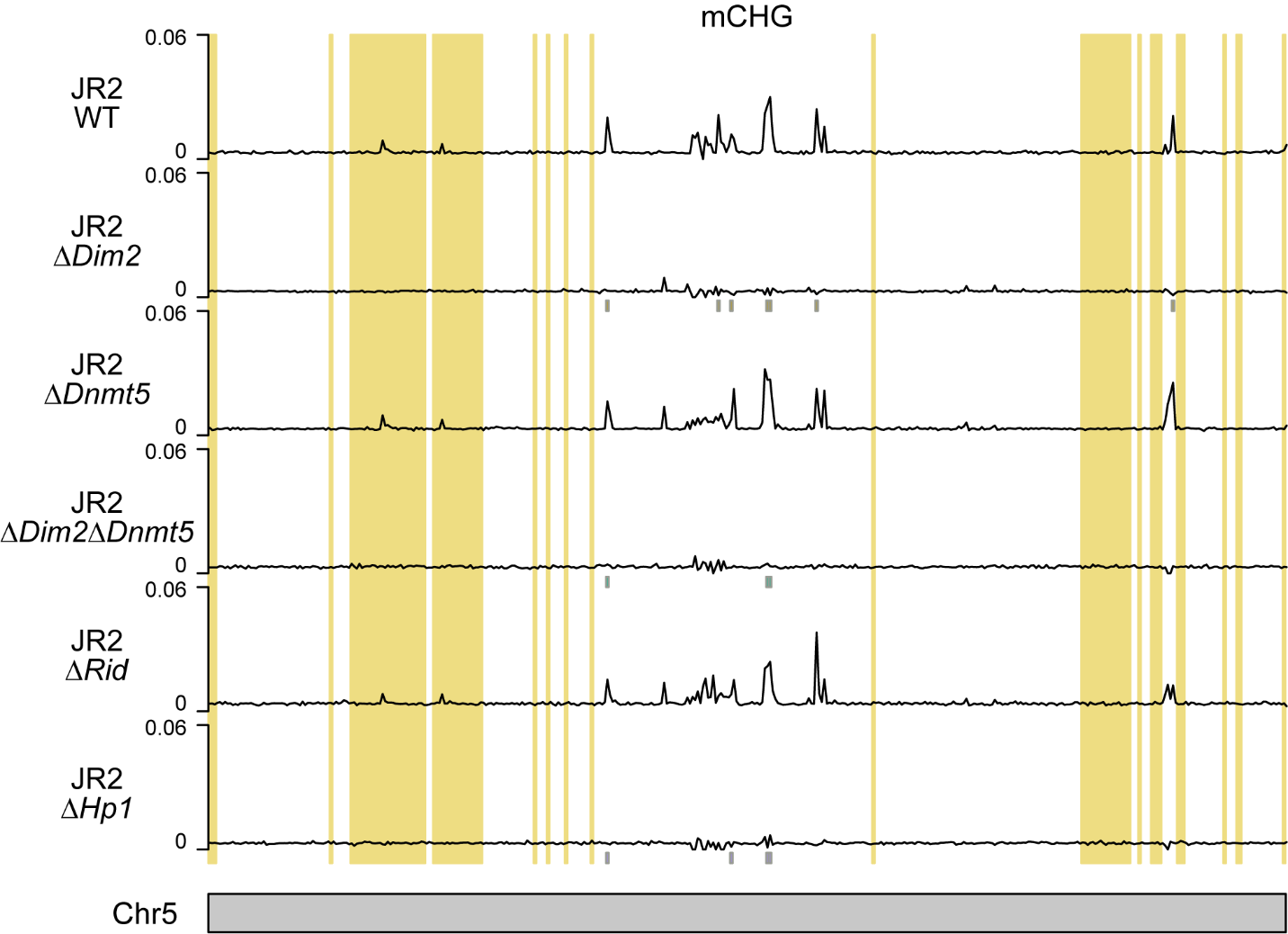

**Figure** **S3. DNA methylation in CHG context.** Whole-chromosome plot displaying the fraction of methylated cytosines for non-overlapping 10 kb windows in CHG context for WT, and DNA methyltransferase and Hp1 deletion mutants with chromosome 5 as an example. Grey boxes, displayed below the DNA methylation tracks, indicate the hypomethylated windows in CHG context from Table 1. Previously defined LS regions (Cook et al., 2020) are highlighted in yellow.

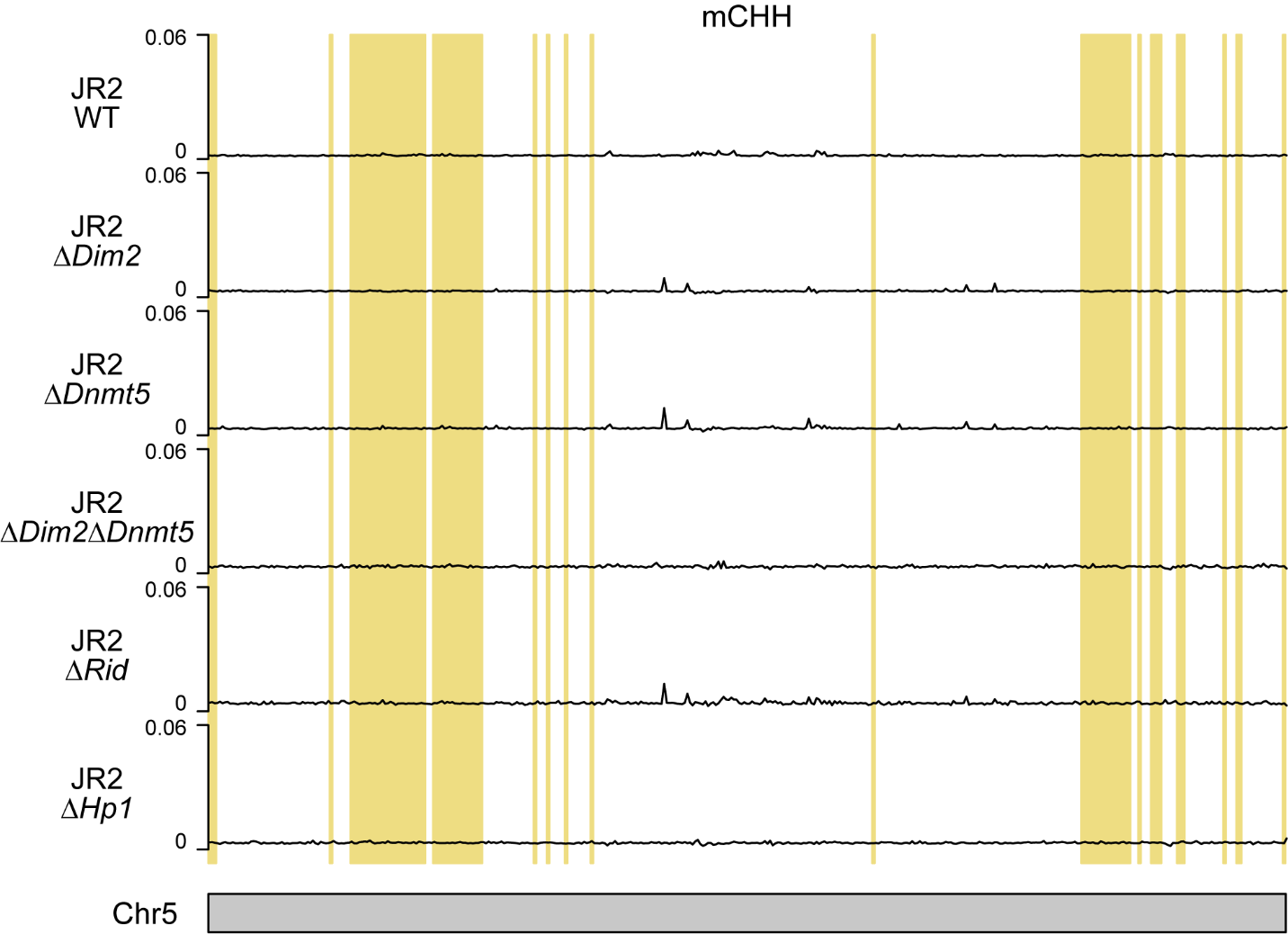

**Figure S4. DNA methylation in CHH context.** Whole-chromosome plot displaying the fraction of methylated cytosines for non-overlapping 10 kb windows in CHH context for WT, and DNA methyltransferase and Hp1 deletion mutants with chromosome 5 as an example. Colored boxes, displayed below the DNA methylation tracks, indicate the hypomethylated windows in CHH context from Table 1. Previously defined LS regions (Cook et al., 2020) are highlighted in yellow.

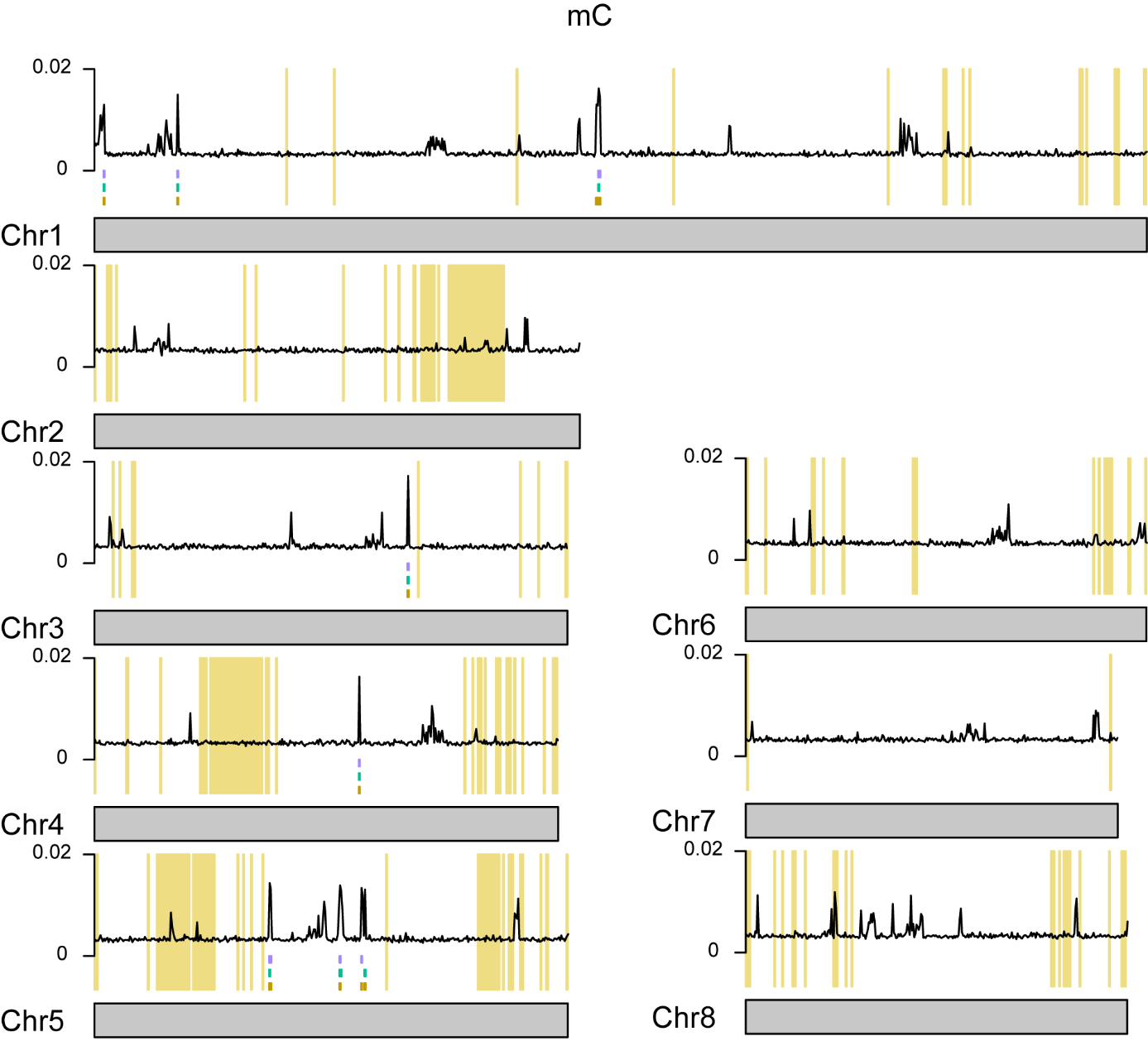

**Figure S5. DNA methylation over the genome.** Whole-chromosome plot displaying the fraction of methylated cytosines for non-overlapping 10 kb windows for wild-type over the whole genome. Colored boxes, displayed below the DNA methylation track indicate the hypomethylated windows from Table 1. Brown ΔDim2, green ΔDnmt5, teal ΔDim2-ΔDnmt5, blue ΔRid, purple ΔHp1. Previously defined LS regions (Cook et al., 2020) are highlighted in yellow.

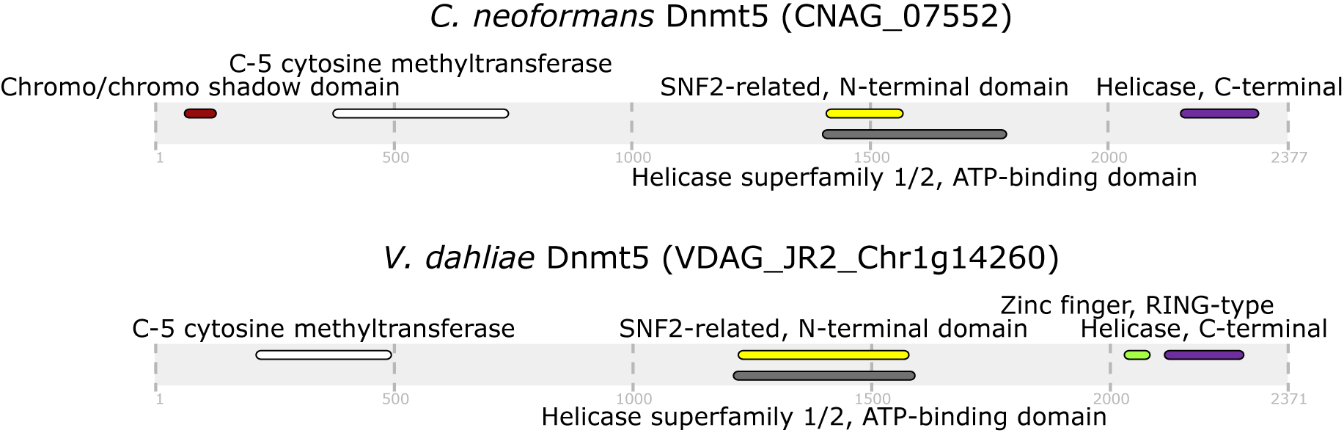

**Figure S6.** Comparison of protein domain structure of C. neoformans and V. dahliae Dnmt5.

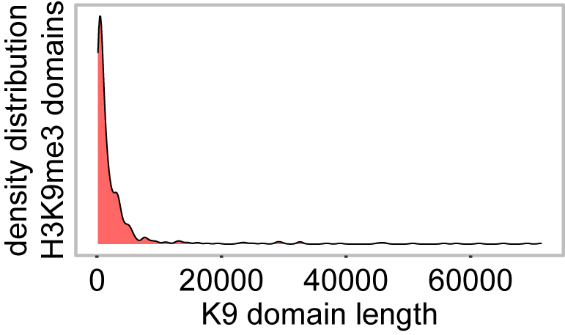

**Figure S7. Distribution of H3K9me3 domain lengths.**

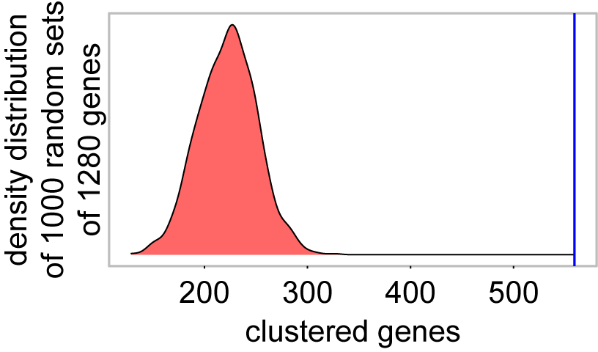

**Figure S8. Genes induced in Hp1 and Dim5 mutants cluster more often than expected based on chance.**

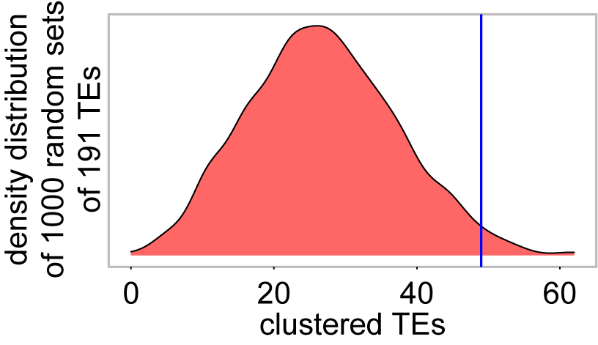

**Figure S9. Transposons induced in Hp1 and Dim5 mutants cluster more often than expected based on chance.**

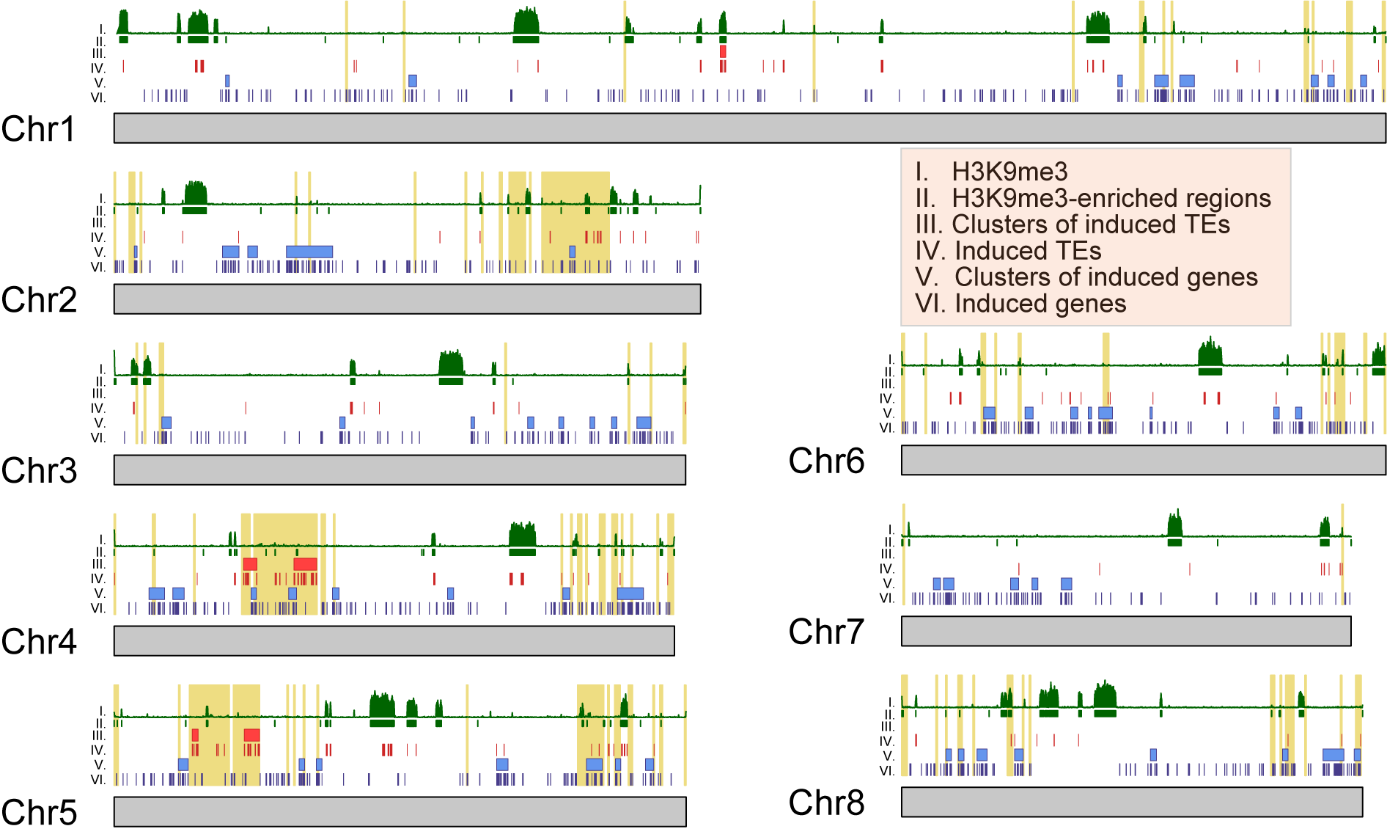

**Figure S10. Clusters of genes and transposons over all chromosomes** Whole-chromosome plots displaying the location of induced genes (in blue) and transposons (in red). Clusters of induced genes and transposons are indicated as blue and red rectangles, respectively. H3K9me3-ChIP signal along the chromosomes is indicated in green in the upper track. Previously defined LS regions (Cook et al., 2020) are highlighted in yellow.
